# Activating words without language: Beta and theta oscillations reflect lexical access and control processes during verbal and non-verbal object recognition tasks

**DOI:** 10.1101/2022.09.21.508907

**Authors:** Francesca M. Branzi, Clara D. Martin, Emmanuel Biau

## Abstract

The intention to name an object modulates neural responses during object recognition tasks. However, the nature of this modulation is still unclear. We established whether a core operation in language, i.e., lexical access, can be observed even when the task does not require language (size-judgment task), and whether response selection in verbal *versus* non-verbal semantic tasks relies on similar neuronal processes. We measured and compared neuronal oscillatory activities and behavioural responses to the same set of pictures of meaningful objects, while the type of task participants had to perform (picture-naming *versus* size-judgment) and the type of stimuli to measure lexical access (cognate *versus* non-cognate) were manipulated. Despite activation of words was facilitated when the task required explicit word-retrieval (picture-naming task), lexical access occurred even without the intention to name the object (non-verbal size-judgment task). Activation of words and response selection were accompanied by beta (25-35 Hz) desynchronisation and theta (3-7 Hz) synchronisation, respectively. These effects were observed in both picture-naming and size-judgment tasks, suggesting that words became activated via similar mechanisms, irrespective of whether the task involves language explicitly. This finding has important implications to understand the link between core linguistic operations and performance in verbal and non-verbal semantic tasks.

## 1. Introduction

Imagine you are at home getting ready to go out. Someone asks you which type of shoes you intend to wear. Your intention to speak will likely activate lexical and phonological representations corresponding to “boots”, because retrieving these representations is crucial to respond to your interlocutor (Strijkers et al. 2012; Branzi et al. 2021). However, what does it happen to those lexical representations if now the task requires to ‘tide up’ and decide whether these boots fit in a shoe box? Would the word representations corresponding to “boots” be activated, even without mandatory and explicit word-retrieval?

Past research has shown that the intention to name an object modulates the neural network and the event-related potential (ERP) responses during object recognition tasks (Strijkers *et al*. 2012; Branzi *et al*. 2021). However, the nature of this modulation is still unclear. The current literature has not yet established whether the intention to name an object facilitates activation of lexical/phonological representations (Strijkers *et al*. 2012), or whether it is a necessary requisite for observing lexicalisation processes (Meyer et al. 1998; Jescheniak et al. 2002; Bles and Jansma 2008; Branzi *et al*. 2021).

Here we investigate whether lexical access takes place even when there is no intention to name (i.e., overtly using language), and whether the neural processes supporting lexical access are quantitatively and/or qualitatively similar to when explicit word-retrieval is intended. Addressing these questions not only is crucial to characterise the neural and cognitive basis of lexical access, a core operation in language, but also to determine whether response selection during semantic tasks requiring verbal *versus* non-verbal responses rely on similar neuro-computations (Fedorenko and Shain 2021).

Neural oscillation patterns are particularly suitable to address these questions as they provide high-temporal resolution to reveal the precise timing of the neural dynamics reflecting spreading activation during lexical access. Furthermore, neuronal oscillation patterns already allowed to establish a link between cognitive and neurophysiological computations in many cognitive domains (Siegel et al. 2012; Friederici and Singer 2015), including language (Meyer 2018; Piai and Zheng 2019), with different oscillatory frequencies being associated to different cognitive processes.

Alpha-beta and theta frequency bands are often found in conceptually-driven word production tasks, such as picture-naming and verb generation (e.g., Ojemann et al. 1989; Piai, Roelofs and Maris 2014; Piai et al. 2015; Piai et al. 2016; Jafarpour et al. 2017; Piai et al. 2017; Piai et al. 2018). Yet, alpha-beta and theta oscillations have been related to different cognitive processes during these tasks. For instance, alpha-beta power desynchronisation (or alpha-beta power decrease) has been related to activation of information within the lexical-semantic system (Piai *et al*. 2015; Roos and Piai 2020). Instead, similarly to the domain of action monitoring (Trujillo and Allen 2007; Cavanagh et al. 2009; Cohen 2011), theta power increases have been associated to cognitive control demands - and especially monitoring control - during the retrieval of lexical representations in language tasks (Geng et al. 2022; Piai and Zheng, 2019).

In the present electroencephalogram (EEG) study, we tested a group of healthy participants and compared their oscillatory activity focusing on alpha-beta and theta frequencies during two different tasks—a picture-naming task and a size-judgment task. Importantly, both picture-naming and size-judgment tasks relied on similar picture processing operations (extraction of visual features, visual-semantic processing for object recognition, response selection) but only one, the picture-naming task, required explicit retrieval of object names. In contrast, the size-judgment task required participants to make a size-judgment providing a manual response to indicate whether an object was “bigger” or “smaller” than a shoebox, but no explicit retrieval of the object name.

Since the retrieval of semantic information has been linked to both alpha-beta desynchronization and theta synchronization irrespective of the intention to speak (e.g., in semantic-based episodic memory tasks; see Piai and Zheng, 2019), we expected that visual object processing would induce an overall significant decrease in alpha-beta power (i.e., desynchronisation) and an increase in theta frequency irrespective of the task. However, since the main goal of the present study was to examine whether the intention to name an object modulated the access to lexical information specifically, we manipulated the ‘cognate status’ of the stimuli in the two tasks (picture-naming and size-judgment).

The cognate status of a word is determined by the extent to which it shares orthographic and phonological features with its translation equivalent in another language. Cognates are translation words that have similar orthographic–phonological forms in two languages (e.g., tomato—English, tomate—Spanish). By contrast, non-cognates are translation equivalents that share only their meaning (e.g., apple—English, manzana—Spanish). Typically, in bilingual and/or multilingual speakers, behavioral and neural differences between non-cognate and cognate processing are observed during picture-naming and indicate lexical/phonological activity (the “cognate effect” (Costa et al. 2000; De Bleser et al. 2003; Costa et al. 2005; Christoffels et al. 2007; Strijkers et al. 2010; Branzi *et al*. 2021). Therefore, as in previous studies, here we tested bilingual participants and we employed the behavioral and neural cognate effect as a proxy for lexical/phonological activity and examined whether this effect (neural and behavioural) varied as a function of the intention to name an object (Strijkers *et al*. 2010; Branzi *et al*. 2021). In fact, since the cognate status of a word is defined by formal overlap and is not correlated with any perceptual or conceptual variable (e.g., Costa *et al*. 2000; De Bleser *et al*. 2003; Costa *et al*. 2005; Christoffels *et al*. 2007; Strijkers *et al*. 2010; Palomar-Garcia et al. 2015), any behavioral or neural difference between non-cognate and cognate processing would reflect a purely lexical/phonological effect.

If a cognate effect were found in both tasks, it would indicate that visual object processing induces automatic activation of lexical/phonological representations, independently of the intention to speak (“spreading activation”, see Dell 1986; Caramazza 1997; Strijkers *et al*. 2012; see also evidence from the picture-word interference studies, i.e., Schriefers et al. 1990; Jescheniak and Schriefers 2001; de Zubicaray et al. 2002; picture-picture interference, e.g., Tipper and Driver 1988; Bles and Jansma 2008; but see other models which do not assume spreading activation in all circumstances, e.g., Levelt 1989; Levelt et al. 1999). If so, enhanced behavioural performance, i.e., faster response times and increased accuracy measures for cognates as compared to non-cognates, should be observed in both picture-naming and size-judgment tasks (Strijkers *et al*. 2010; Branzi et al. 2020; Branzi *et al*. 2021).

Despite a lack of evidence, the literature indicates that retrieving lexical information from memory is associated with power decreases in the alpha-beta band, similarly to the episodic-memory domain (Piai and Zheng, 2019). Since lexical retrieval depends on the activation level of the target lexical representations (Dell 1986; Levelt *et al*. 1999; Finkbeiner et al. 2006), a greater desynchronisation of neural alpha-beta oscillations should be observed for the less-activated representations, i.e., for non-cognates as compared to cognates. Note that if this neural cognate effect were found in both tasks, it would indicate that lexical access during object naming and object-size categorization reflects the same spreading activation mechanism. Still, the neural cognate effect might be observed in an earlier time window for the picture-naming task as compared to the size-judgment task (Strijkers *et al*. 2012), which would indicate similar processes across tasks, but faster when explicit word-retrieval is required.

Furthermore, if theta power reflects monitoring control processes during response selection (Cavanagh and Frank 2014; Cohen 2014; Geng *et al*. 2022), this effect should be stronger in the picture-naming task as compared to the size-judgment task for cognate stimuli only. In fact, in bilingual speakers the intention to name an object (picture-naming task) should induce faster dual-language activation (Strijkers *et al*. 2012). This should affect cognates only because cognate, but not non-cognate words, receive activation from both the target (e.g., *tomato* in English) and non-target languages (translation equivalent, e.g., *tomate* in Spanish). This simultaneous activation might increase conflict between target and non-target translation equivalents that share lexical/phonological representations and therefore the need of monitoring control (Li and Gollan 2018). In accord with the view that theta oscillations synchronise when conflict between representations increases (Cavanagh and Frank 2014; Cohen 2014; Geng *et al*. 2022), theta power responses should increase during the picture-naming task as compared to the size-judgment task for cognate stimuli only.

Finally, we investigated whether the activation of knowledge hypothetically indexed by alpha-beta activity here (i.e., semantic and lexical activation) interacted with processes supported by theta activity (i.e., monitoring control) during lexical access. To evaluate the functional relationship between the theta phase and the beta amplitude during the two tasks, we conducted cross-frequency coupling analysis. Despite it is reasonable to hypothesise that the intention to name an object would modulate interactions between two different oscillations reflecting distinct cognitive processes during lexical access, the exact pattern of results is hard to predict from the current literature. To our knowledge, in fact, our is the first study to examine cross-frequency interactions in a task measuring lexical-semantic activation. It is possible that, similarly to the expected theta power effects, the interplay between theta and beta oscillatory responses might be modulated by increased control demands during response selection. This hypothesis would be in accord with results from studies in working memory research that have shown increase of beta-theta coupling proportional to increase in control demands (Daume, Graetz, et al. 2017; Daume, Gruber, et al. 2017).

## 2. Materials and Methods

### Participants

Twenty early and high-proficient Catalan/Spanish bilinguals took part in the experiment and received monetary remuneration for their participation (25 females; mean age: 21.3 years ± 1.9; for further details, see Branzi et al. 2014). Half of the participants were dominant in Spanish whereas others were more dominant in Catalan. However, all participants were early bilinguals, equally proficient in both languages (see self-assessed proficiency scores are reported in (Branzi *et al*. 2014). All participants were right-handed and had normal or corrected-to-normal vision. The data set of two participants was excluded from the analyses due to excessive movement artifact contamination. The analyses were applied on the remaining eighteen participants. The study was approved by the local ethics committee. Written informed consent was obtained from all participants.

### Stimuli

One hundred and twenty-eight line-drawings of objects were selected from different databases (Snodgrass and Vanderwart 1980; Szekely et al. 2004). The pictures represented objects belonging to a wide range of semantic categories (e.g., animals, body parts, buildings, furniture). In both the picture-naming and size judgment tasks, the objects had to be named in Spanish and Catalan and could refer to either cognate or non-cognate words. The cognate status of the corresponding names was controlled to present fifty per cent of the items as Spanish/Catalan cognates and the other fifty per cent as non-cognates. The mean lexical frequency (LEXESP; Sebastián-Gallés et al. 2000) of the picture names was balanced between cognates and non-cognates (non-cognates: 1.03 ± 0.6; cognates: 1.14 ± 0.6; *t* (126) = -1.075, *p* = .284). Pictures were grouped together in order to create four experimental lists, which were then randomized across participants. Even if each participant was not presented with the very same pictures during the picture-naming and size-judgment tasks (see below), the very same stimuli and lists were employed for the two tasks across participants.

### Experimental procedure: Picture-naming and size-judgment tasks

After having filled in the informed written consent and a language use/proficiency questionnaire, participants were tested individually in a soundproof room. Written instructions were presented in their native language (L1). Participants performed a picture-naming task and a size-judgment task in their L1^1^ during a single session. In both picture-naming and size-judgment tasks participants were presented with a set of 64 pictures. Trial order within the block was randomized. During the picture-naming task participants were required to name pictures depicting concrete objects in their L1. Instead, in the size-judgment task participants performed a size-judgment task on the objects depicted in the pictures (“Is this object bigger/smaller than a shoebox?”; see Dobbins et al. 2004; Branzi et al. 2016; Branzi *et al*. 2021). In this two-alternative forced choice setting, participants provided a yes/no response via button press. Note that participants were told that to perform the size-judgement task correctly they had to consider the size of the object in the real world. Despite the two tasks differed substantially in terms of type of response given, they both required to access to some semantic knowledge. Participants were not familiarized with the picture names beforehand to avoid repetition priming effects (e.g., Guo et al. 2011; Misra et al. 2012).

The stimuli were presented using Presentation software (Neurobehavioral systems: http://www.neurobs.com/). Vocal response latencies (picture-naming task) and button press responses (size-judgment task) were recorded from the onset of the stimuli. In both tasks, each trial began with a blank screen for 1000 ms, the picture appeared for 1500 ms at the centre of the screen on a black background. Then, a fixation cross was presented for 500 ms.

### Behavioral data and statistical analyses

The experiment used a full within-subject design. The mean correct response rates (i.e., accuracy) and the mean reaction times of the correct trials comprised between mean reaction times ± two standard deviations range were computed in the four conditions, i.e., picture-naming non-cognate (NamingNC), picture-naming cognate (NamingC), size-judgment non-cognate (Size-judgmentNC) and size-judgment cognate (Size-judgmentC), separately for each participant. To compare the cognate effect across the two tasks we used a two-way repeated-measure Analyses of Variance (ANOVAs) with the within-subjects factors Task (picture-naming *versus* size-judgment) and Cognate status (non-cognate *versus* cognate). Statistically significant interactions were assessed via planned post-hoc *t*-tests. For the pairwise comparisons we also provided an effect size value (Cohen’s *d*) and a Bayes factor value (BF10 > 3 suggests substantial evidence for a difference between the pairs and BF10 < 0.3 suggests substantial evidence for a null effect, see Jeffreys 1961). Reporting Bayes factors is useful for hypothesis testing because they provide a coherent approach to determining whether non-significant results support the null hypothesis over a theory, or whether the data are just insensitive.

#### EEG recording and preprocessing

Electrophysiological data was recorded (Brain Vision Recorder 1.05; Brain Products) from 38 tin electrodes placed according to the 10–20 convention system. An electrode placed on the tip of the nose was used as a reference. Two bipolar electrodes were placed next to and above the right eye to register ocular movements. Electrode impedances were kept below 5 kΩ and EEG signal was recorded with a high cut-off filter of 200 Hz, with a sampling rate of 500 Hz. Offline EEG pre-processing involved EEG data being pre-processed offline using Fieldtrip (Oostenveld et al. 2011). Continuous EEG signals were bandpass filtered (standard non-causal two-pass Butterworth filters) between 0.1 Hz and 100 Hz and bandstop filtered (48-52 Hz and 98-102 Hz) to remove line noise at 50 and 100 Hz. Data were epoched from 1000 ms before the stimulus onset to 1000 ms after stimulus onset. Trials and channels with artefacts were excluded by visual inspection before applying an independent component analysis (ICA) to remove components related to ocular artefacts. Excluded channels were then interpolated using the method of triangulation of nearest. After re-referencing the data to an average reference across all electrodes, the remaining trials with artefacts were manually rejected by a final visual inspection (on average, 6.77 ± 4.37 trials per participant (i.e., 5.30 ± 3.41 per cent of total trials per participant).

#### EEG data processing and statistical analyses

Only the correct trials were included in the statistical analysis. Time-frequency decomposition was conducted to each electrode using a Morlet wavelet (width: 5 cycles, from 1 to 40 Hz; 1 Hz step and 20 ms time steps) and frequency analyses were performed for each trial in the four conditions, i.e., NamingNC, NamingC, Size-judgmentNC and Size-judgmentC, separately for each participant. Further, background fractal activity was attenuated in the TFR by subtracting 1/f characteristic from the spectral power using an iterative linear fitting procedure (Griffiths et al. 2021). This step generated two vectors: one vector contained the values of each wavelet frequency A, while the other vector contained the power spectrum for each electrode-sample pair B. Both vectors were then put into log-space to provide a linear line to get the slope and intercept of the 1/f curve. The linear equation Ax = B was resolved using least-squares regression, where x is an unknown constant describing the curvature of the 1/f characteristic. The 1/f fit Ax was then subtracted from the log-transformed power spectrum B. The corrected power was then averaged across trials in separate conditions for each participant.

The differences of power between the two contrasts NamingNC - NamingC and Size-judgmentNC – NamingC were first statistically assessed by applying dependent *t*-tests using Monte-Carlo cluster-based permutation tests (Maris and Oostenveld 2007) with an alpha cluster-forming threshold set at 0.05, three minimum neighbour channels, 2000 iterations, and cluster selection based on maximum size. Cluster-based permutation statistics were applied on the mean beta power (25-35 Hz) averaged across the time windows of interest determined in the contrasts NamingNC *versus* NamingC and Size-judgmentNC *versus* Size-judgmentC. Further, the normalised mean beta power relative to a baseline preceding the stimulus onset (-700 to -200 ms with respect to stimulus onset) was averaged across the electrodes of the regions of interests in the 25-35 Hz frequency band for the four conditions separately, and exported for statistical assessments. First, the mean of normalized beta power was compared to zero in the four conditions by applying one-sample *t*-tests (two-tailed) to confirm a significant decrease of beta activity during stimulus processing. The *p*-values were corrected for multiple comparisons using the Bonferroni correction (α = .025). Second, the differences of beta power between conditions were statistically assessed by means of a two-way repeated-measure ANOVA with Task (picture-naming *versus* size-judgment) and Cognate Status (non-cognate *versus* cognate) as within-subjects factors. Statistically significant interactions were further assessed via planned post-hoc *t*-tests. For the pairwise comparisons we also provided an effect size value (Cohen’s *d*) and a Bayes factor value (Jeffreys 1961). The same procedure (i.e., extraction of normalized power and statistical analyses) was applied for the analysis of 3-7 Hz theta and 8-12 Hz alpha bands. To anticipate the results, we did not find any significant effect of alpha activity during picture-object processing, across all conditions (see **Figure S1**). Thus, the following planned analyses were restricted to beta and theta frequencies.

Finally, we performed a searchlight-based analysis to contrast the amplitude of the cognate effect on beta power in the cluster of interest against all the remaining electrodes of the scalp. First, we iterated through each outside electrode randomly. Second, we identified its immediate neighbours to create a mini cluster for each iteration, i.e., nearest electrodes to the iteration electrode in the 2-dimensional space. For each iteration, the size of the mini cluster was of 11 neighbour electrodes in the picture-naming task, and of 8 neighbour electrodes in the size-judgment task. Third, we computed the mean amplitude of the beta power in the non-cognate and cognate conditions within this mini cluster. Fourth, the mean value was contrasted between the non-cognate and cognate conditions for each mini-cluster, and the resulting t-statistic was added to a distribution describing the amplitude of the cognate effect on beta power across the scalp, for the picture-naming and size-judgment tasks separately. Fifth, the *p*-value was derived by comparing the cognate effect in the regions of interest to the scalp distribution with a permutation test (2000 permutations). Therefore, this approach allowed to infer the extent to which the cognate effect observed in the regions of interest effectively deviated from the rest of the scalp.

#### Theta-beta phase-amplitude coupling (PAC) and statistical analyses

In this analysis we did not consider the alpha frequency band because, as reported above, we did not find any significant effect of alpha activity during picture-object processing, across all the conditions (*see* **Figure S1** for further analysis on the alpha band). Instead, we investigated the functional relationship between theta and beta oscillations and particularly whether theta-beta coupling would reflect a mechanism for the collaborative functioning of lexical-semantic activation (beta effect) and response monitoring (theta effect) during response selection. We performed PAC analyses directly applied on the time-windows and regions of interest determined in the previous EEG analysis step. We expected the strength of theta-beta coupling to be modulated by increase of control demands for cognates during the picture naming task *versus* the size-judgment task.

To assess the extent to which the beta activity coupled to theta phase, we calculated the modulation index (MI) from the correct trials only (Tort et al. 2010; Griffiths *et al*. 2021; Biau et al. 2022). First, the peaks in the theta and beta frequency bands were calculated by estimating power across all electrodes from the regions of interest in the picture-naming task (NamingNC and NamingC) and the size-judgment task (Size-judgmentNC and Size-judgmentC), with the same time-frequency decomposition method as above. The most prominent peaks in the theta (3-7 Hz) and beta (13-30 Hz) bands measured during picture-naming and size-judgment tasks were extracted for each individual participant using the Matlab function ‘findpeaks’. In the picture-naming task, the mean theta peak across participants was found at 5.04 ± 0.13 Hz and the mean beta peak was found at 29.84 ± 0.79 Hz. In the size-judgment task, the mean theta peak across participants was found at 5 ± 0.13 Hz and the mean beta peak was found at 29.59 ± 0.65 Hz. To obtain an equal number of trials across conditions before the MI calculation, the same number of trials across all conditions was determined by taking 80% of the smallest number of available correct trials across all the conditions (average minimum number of trials: 22.61 ± 3.59). The 80% subsampling was done to ensure that some participants were not overrepresented in the resampling procedure due to using 100% of their available data, as well as to vary the set of trials in the condition determining the minimum number of trials across iterations (Keitel et al. 2018). Second, the time-series of the electrodes of interest were duplicated and filtered separately: the first time-series was filtered around the theta peak (± 0.5 Hz) and the second time-series was filtered around the beta peak (± 5 Hz). Third, the Hilbert transform was applied to the theta and beta filtered time-series to extract the phase of the former and the power of the latter. Fourth, beta power was binned into 12 equidistant bins of 30° according to the theta phase. The binning was computed for each trial and electrode separately. The MI was computed by comparing the observed distribution to a uniform distribution for each trial and electrode. The MI was then averaged across the trials and electrodes in each condition separately for statistical assessments. That is, first the mean PAC was compared to zero in the four conditions by applying one-sample *t*-tests (two-tailed) to confirm that beta activity coupled to theta phase during visual object processing. The *p*-values were corrected for multiple comparisons using the Bonferroni correction (α = .025). Then, the statistical differences of mean PAC across conditions were assessed via two-way repeated-measure ANOVA with Task (picture-naming *versus* size-judgment) and Cognate Status (non-cognate *versus* cognate) as two within-subject factors. Statistically significant interactions were further assessed via planned post-hoc *t*-tests.

#### Data and scripts availability statement

Data and scripts to reproduce the results reported in this manuscript will be made available upon publication of the manuscript. Further information or requests should be directed to the corresponding authors.

## 3. Results

### Behavioural results

The effect of “intention to name an object” on lexical access was assessed by comparing the cognate effect across the two tasks. The results relative to accuracy measures (proportion of correct responses) are reported in **Figure 1**: NamingNC: 0.72 ± 0.12; NamingC: 0.84 ± 0.09; Size-judgmentNC: 0.79 ± 0.10 and Size-judgmentC: 0.85 ± 0.09. The ANOVA’s results revealed a significant main effect of Cognate Status, suggesting that retrieving cognate representations is easier than retrieving non-cognate representations [F (1,17) = 30.22; *p* < .001; η*p2* = .64]. Although suggesting a tendency towards significance, the main effect of Task [F (1,17) = 3.12; *p* = .09; η*p*2 = .16] and the interaction between Task and Cognate Status were not significant [F (1,17) = 3.52; *p* = .08; η*p*2 = 0.17].

**Figure 1.**
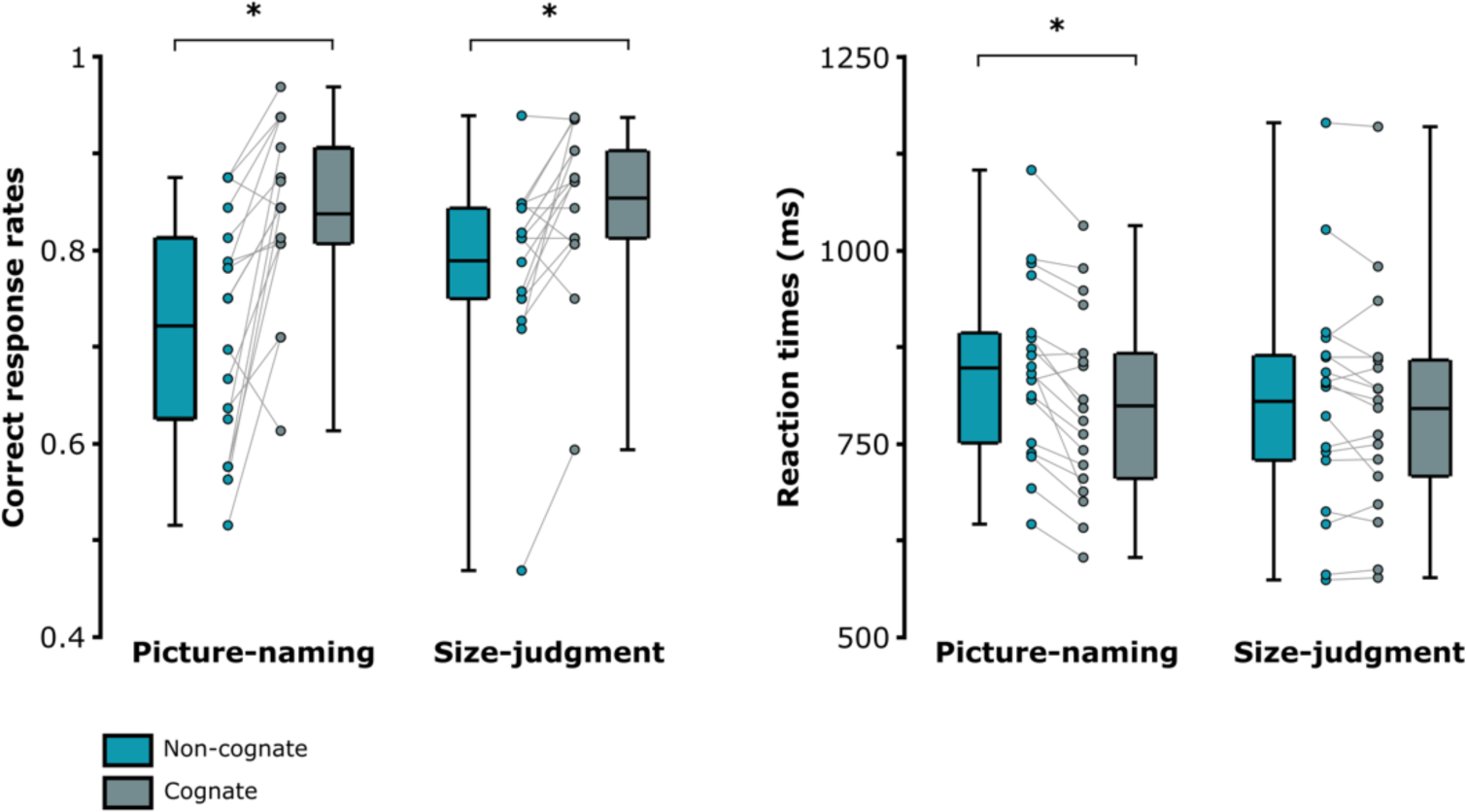
Behavioural results. Boxplots reflect accuracy (proportion of correct responses) and response times (ms) in the picture-naming and size-judgment tasks. Significant differences are evidenced with black stars. The bottom part of the boxplots indicates the first quarter of the score distributions, the upper part of the boxplots indicates the third quarter of the score distributions. The error bars indicate the minimum and maximum scores from the distributions.

The mean reaction times across conditions are depicted in **Figure 1**: NamingNC: 848 ± 115 ms; NamingC: 799 ± 120 ms; Size-judgmentNC: 805 ± 147 ms and Size-judgmentC: 795 ± 142 ms. In line with accuracy results, ANOVA’s results revealed a significant main effect of Cognate Status, suggesting that responses for non-cognates were overall slower as compared to those for cognates [F (1,17) = 25.153; *p* < .001; η*p2* = .597]. The main effect of Task was not significant [F (1,17) = 0.97, *p* = .338; η*p2* = .054]. However, the interaction between Task and Cognate Status was significant [F (1,17) = 11.814; *p* = .003; η*p2* = .41]. Planned comparisons revealed that the cognate effect, i.e., cognate conditions being faster as compared to non-cognate conditions, was observed in the picture-naming task only [Naming: *t* (17) = -5.366; *p* < .001 (adjusted), Cohen’s *d* = -1.265, BF10 = 1020.672; Size-Judgement: *t* (17) = -1.286; *p* = .216 (adjusted); Cohen’s *d* = -.303, BF10 = .869].

To summarise, behavioural results suggest that the “intention to name an object” is not mandatory to observe lexical/phonological access. However, the cognate effect modulated both response times and accuracy only during the picture-naming task. Therefore, these results suggest that “intention to name an object” might modulate the strength of lexical/phonological activity (cognate effect).

### Picture-object processing and lexical access lead to modulations in beta (but not alpha) frequency band

The effect of “intention to name an object” on lexicalization processes was assessed by comparing the time-frequency representations (TFRs) between non-cognate *versus* cognate trials in the picture-naming and size-judgment tasks. This analytic step served to determine the time-window of interest containing the cognate effect indexing lexical access through oscillatory response modulations. Firstly, the TFRs in **Figure 2a** did not suggest any difference of power in the 8-12 Hz alpha band (*see* **Figure S1**). In contrast, the TFRs suggested a strong decrease of power in the expected beta band (25-35 Hz) when comparing non-cognate *versus* cognate conditions in both tasks (i.e., cognate effect; **Figure 2a**). The cognate effect was observed between 160 to 260 ms after the stimulus onset in the picture-naming task, a similar effect was observed later in the size-judgment task, i.e., between 260 to 380 ms after stimulus onset. The cluster-based analysis in these specific time-windows confirmed a significant decrease of beta power (25-35 Hz) in the cognate as compared to the non-cognate condition, in both the picture-naming [time-window: 160-260 ms; *p* < .001, positive cluster size = 26.94; mean *t*-statistic within cluster = 2.45] and the size-judgment task [time-window: 260-380 ms; *p* < .001, positive cluster size = 18.44; mean *t*-statistic within cluster = 2.30]. No significant negative cluster was found. **Figure 2b** depicts the topographies relative to beta desynchronisation for the non-cognate *versus* cognate contrast for the two tasks separately. These results reveal that the cognate effect is reflected by beta desynchronisation in similar regions of the scalp (centro-parietal scalp region) in the picture-naming and size-judgment tasks.

**Figure 2.**
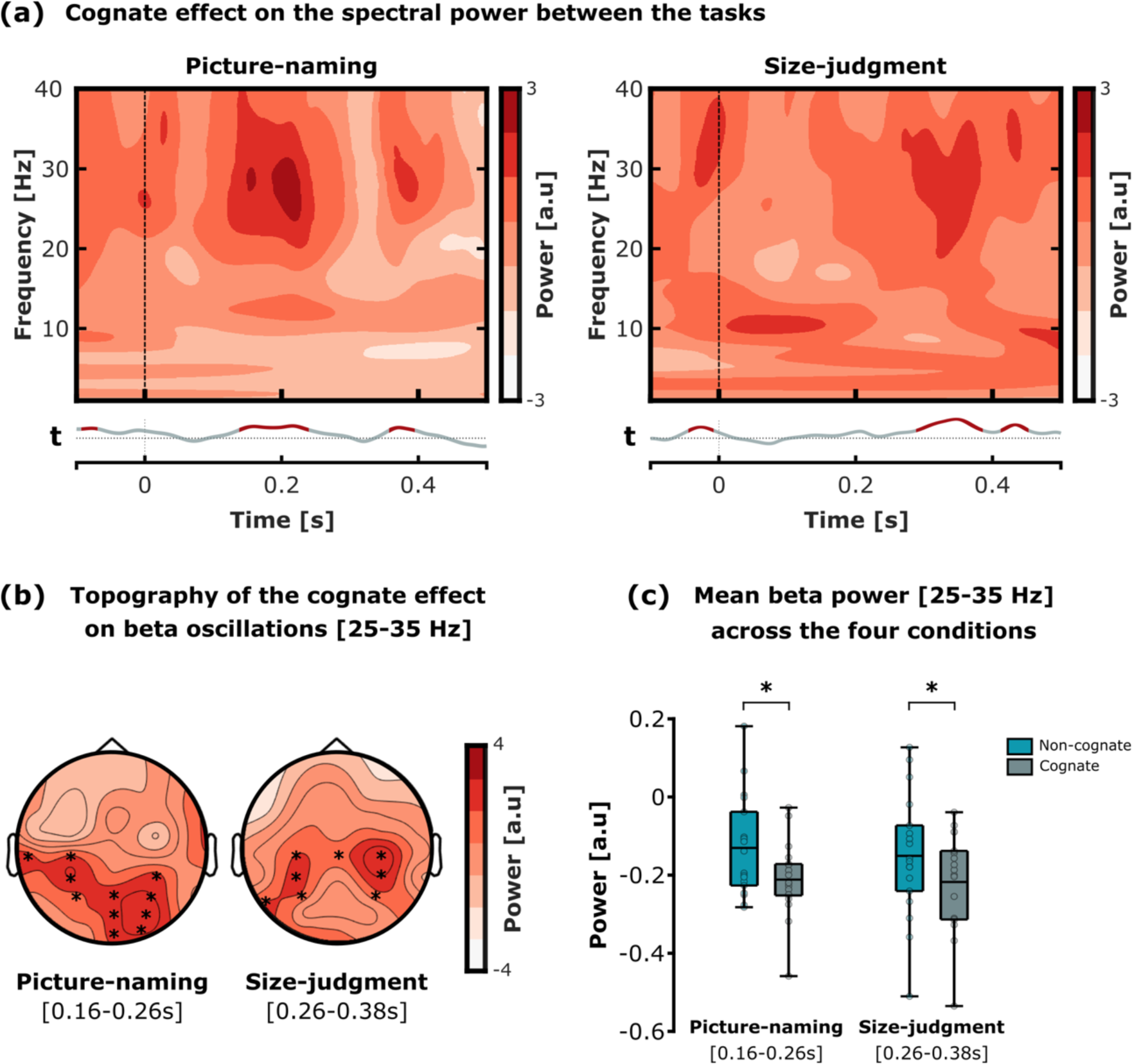
Cognate effect (non-cognates *versus* cognates) modulation on beta oscillations during the picture-naming and size-judgment tasks. (a) Time-frequency representations (TFRs) of the spectral power difference, i.e., non-cognate *versus* cognate, in the picture-naming (left) and in the size-judgment task (right). The TFRs depict the average of all electrodes included in the significant clusters, evidenced with black stars in the topoplots of panel b. The line below the two TFRs represents the *t*-values from the statistical comparisons of the mean 25-35 Hz beta power between the non-cognate *versus* cognate condition for each time-point (the time-points with a significant difference are evidenced in red; *p*-values cluster-corrected for multiple comparisons). (b) Topographies of the difference of beta power between the non-cognate *versus* cognate condition in the picture-naming (left) and size-judgment task (right). The time-windows of interest for the picture-naming and size-judgment task were respectively 160 to 260 ms and 260 to 380 ms after the stimulus presentation onset. The electrodes included in the significant clusters are evidenced with black stars. (c) Boxplots of the mean beta power (25-35 Hz) across the cognate and non-cognate conditions and tasks (significant differences are evidenced with black stars). The time-window of interest for the picture-naming and size-judgment task were respectively 160 to 260 ms and 260 to 380 ms after the stimulus presentation onset. The bottom part of the boxplots indicates the first quarter of the score distributions, the upper part of the boxplots indicates the third quarter of the score distributions. The error bars indicate the minimum and maximum scores from the distributions.

We further examined whether picture-object processing induced significant beta desynchronization across all conditions (see Piai and Zheng, 2019). First, the mean normalized beta power was computed for all conditions (correct trials only) across the significant electrodes and in the time-windows of interest (*see* **Figure 2**). The results for the picture-naming task (significant electrodes: T3, C3, CP3, CP4, P3, P4, Pz, POz, PO2, Oz and O2; time-window: 160-260 ms) and the size-judgment task (significant electrodes: T5, Cz, C3, C4, CP3, CP4, P3 and P4; time-window: 260-380 ms) are reported in **Figure 2c**. Then, four one-sample *t*-tests revealed a significant decrease of mean beta power in response to the stimulus presentation across all conditions [NamingNC: -0.13 ± 0.13, *t* (17) = -4.278, *p* < .001, Cohen’s *d* = -1.008, BF10 = 67.324; NamingC: -0.21 ± 0.1, *t* (17) = -9.318, *p* < .001, Cohen’s *d* = -2.196, BF10 = 318049; Size-judgmentNC: -0.15 ± 0.16, *t* (17) = -4.01, *p* < .001, Cohen’s *d* = -.945, BF10 = 40.68; Size-judgmentC: -0.22 ± 0.13, *t* (17) = -7.346, *p* < .001, Cohen’s *d* = -1.731, BF10 = 15785.3; Significant *p*-values were corrected for multiple comparisons at α = .0125].

In line with our hypothesis, these results confirm that picture-object processing induced significant beta desynchronisation independently from the task and condition. The ANOVA’s result revealed a significant effect of Cognate status [F (1,17) = 12.56; *p* < .001, η*p2* = .43], establishing a greater beta desynchronisation for cognates as compared to non-cognates. However, no significant effect of Task [F (1,17) = 0.1, *p* = .75, η*p*2 < .001] or interaction between Task and Cognate status [F (1,17) = 0.14, *p* = .71, η*p*2 < .001] was found. These results confirmed greater desynchronisation of the beta oscillations when participants processed cognate as compared to non-cognate conditions, independently from the task. Further, the searchlight-based analysis revealed that the amplitude of the cognate effect on beta power in the regions of interest was significantly greater than any other size-matching searchlight-based regions over the scalp (*p* < .001 in both the picture-naming and the size-judgment tasks; *see* **Figure S3** for histogram of searchlight statistics). To control for potential confounds driven by the difference of electrode pool sizes, we performed the exact same time-frequency decomposition of the spectral power difference with a common pool of electrodes overlapping the two clusters of interest reported in the picture-naming and size-judgment tasks. For this control analysis, the common cluster of interest contained the following electrodes: C3, CP3, P3, C4, CP4, P4, Cz, CPz, and Pz. Results replicated the present results, with the processing of picture-objects inducing a significant beta desynchronisation across tasks and conditions. Furthermore, the cognate conditions induced a greater beta desynchronisation of the beta oscillations as compared to non-cognate conditions in both tasks (*see* **Figure S2**). Crucially, we verified that the timing difference relative to the cognate effect (beta desynchronization) observed between the picture-naming (160 to 260 ms with respect to the stimulus onset) and the size-judgment tasks (260 to 380 ms with respect to the stimulus onset) could not reflect only differences in the type of (verbal *versus* manual) response preparation processes. To this end, we assessed the difference of power spectrum between the picture-naming and size-judgment tasks, after collapsing the cognate and non-cognate trials together (*see* **Figure S5a**). The resulting TFR showed that there was no difference of power in the low frequencies of interest (< 35 Hz) within the time-window containing the cognate effect (i.e., approximatively from 100 to 400 ms with respect to the stimulus onset).

Finally, since we found that the cognate status modulated both behavioral and beta oscillatory responses, we tested whether the magnitude of beta desynchronisation induced by the cognate effect predicted the magnitude of the behavioural cognate effect (i.e., accuracy measures). We did so by conducting Pearson’s correlation analyses for the two tasks separately. Results did not reveal any significant correlation between behavioral performance and beta responses for the non-cognate *versus* cognate contrast (*see* **Figure S4 and S5b)**.

Together, the key results from this analysis revealed that (1) picture-object recognition and lexical/phonological activity modulated neural responses in the beta-frequency band (**Figures 2a and 2c**); (2) beta-band neural effects occur earlier when the task requires explicit word-retrieval, in line with previous studies (Strijkers *et al*. 2012). However, (3) the strength of lexical/phonological activity reflected by beta (magnitude of the cognate effect) seems to be independent from whether the task requires explicit retrieval of the object name (**Figure 2c**). Finally, (4) the beta-band neural effects reflecting lexical/phonological activity showed similar scalp distribution in the two tasks, i.e., parietal and occipital electrodes, in line with previous reports (Strijkers *et al*. 2012).

Picture-object processing and increased monitoring demands during picture-naming modulate theta activity.

We examined whether picture-object processing induced a significant increase of theta power reflecting monitoring processes during response selection independently from the condition or the task (for a review, see Piai and Zheng, 2019). We applied this analysis directly in the two time-windows of interest determined above (i.e., exhibiting the greater cognate effect on beta oscillation responses in the two tasks). In line with our hypothesis, we found that theta activity (3-7 Hz) increased during picture-object processing as compared to pre-stimulus baseline (**Figure 3a, left panel**). Four one-sample *t*-tests confirmed a significant increase of theta activity during picture-object processing across all conditions and tasks [NamingNC: 0.47 ± 0.23, *t* (17) = 8.93, *p* = < .001, Cohen’s *d* = 2.104, BF10 = 181214.807; NamingC: 0.55 ± 0.34, *t* (17) = 6.84, *p* < .001, Cohen’s *d* = 1.613, BF10 = 6869.73; Size-judgmentNC: 0.46 ± 0.27, *t* (17) = 7.19, *p* < .001, Cohen’s *d* = 1.695, BF10 = 12298.681; Size-judgmentC: 0.36 ± 0.22, *t* (17) = 6.95, *p* < .001, Cohen’s *d* = 1.639, BF10 = 8271.203; Significant *p*-values were corrected for multiple comparisons at α = .0125].

**Figure 3.**
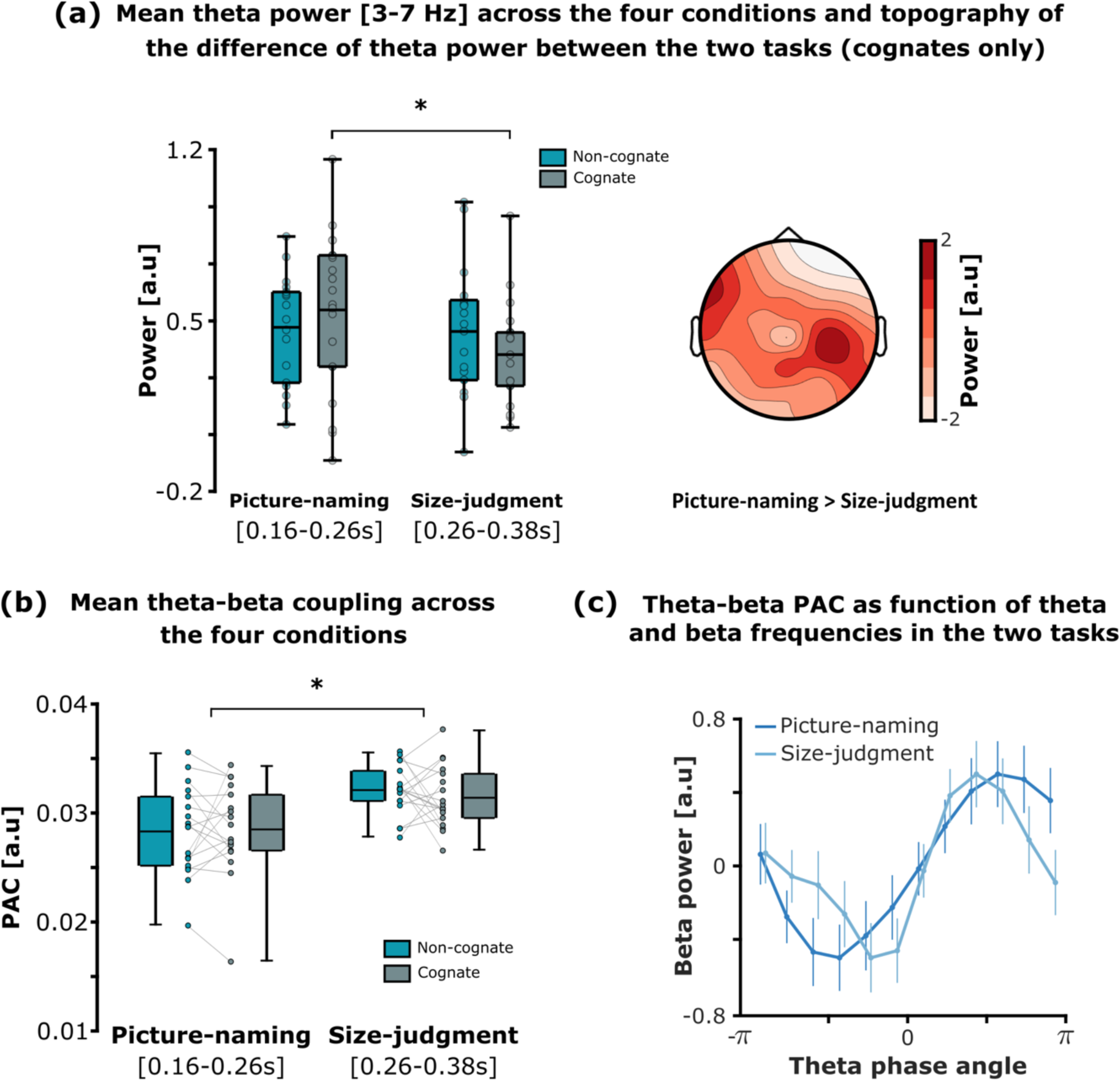
Cognate effect (non-cognates *versus* cognates) modulation on theta power (3-7Hz) and theta-beta phase-amplitude coupling (PAC) in the picture-naming and size-judgment tasks. (a. left) Boxplots of the mean theta power (3-7 Hz) across the cognate and non-cognate conditions and tasks. The time-window of interest for the picture-naming and size-judgment task were respectively 160 to 260 ms and 260 to 380 ms after the stimulus presentation onset. The bottom part of the boxplots indicates the first quarter of the score distributions, the upper part of the boxplots indicates the third quarter of the score distributions. The error bars indicate the minimum and maximum scores from the distributions. (a. right) Topography of the difference of theta power (picture-naming *versus* size-judgment task) in the cognate condition only. No significant cluster was found, reflecting only a tendency to differ as depicted in the left boxplots (i.e., difference between the grey boxes). (b) Boxplots of the mean theta-beta PAC across the cognate and non-cognate conditions and tasks (significant differences evidenced with black stars). The time-windows of interest for the picture-naming and size-judgment task were respectively 160 to 260 ms and 260 to 380 ms after the stimulus presentation onset. The bottom part of the boxplots indicates the first quarter of the score distributions, the upper part of the boxplots indicates the third quarter of the score distributions. The error bars indicate the minimum and maximum scores from the distributions. (c) Beta power as a function of theta phase in the regions of interest, for the picture-naming and size-judgment task (non-cognate and cognate trials collapsed together within task). The fluctuation of beta power (y-axis) during the size-judgment task appears to concentrate more towards the theta phase (0° on the x-axis) as compared to the picture-naming task. Although this last analysis provides visual support only, results suggest that more narrow-band oscillations account for theta-beta PAC in the size-judgment task as compared to the picture-naming task.

A two-way repeated-measure ANOVA with Task (picture-naming *versus* size-judgment) and Cognate status (non-cognate *versus* cognate) as within-subject factors revealed a significant interaction [F (1,17) = 5.609; *p = .*03; η*p2* = .248]. Planned pairwise comparisons revealed increase of theta power in the picture naming task *versus* the size-judgment task for cognates [*t* (17) = 2.341, *p* = .032 (adjusted), Cohen’s *d* = 0.552, BF10 = 4.048], but not for non-cognates [*t* (17) = 0.272, *p* = 0.788 (adjusted), Cohen’s *d* = .064, BF10 = 0.302). The main effect of Task [F (1,17) = 2.677, *p* = .12, η*p2* = .136] and Cognate Status [F (1,17) = 0.094; *p* = .762, η*p2* = .006] were not significant. Although no electrode showed significant effects, the topoplot (**Figure 3a**, right panel) suggested that the difference of theta power in the cognate condition between the picture-naming and the size-judgment tasks peaks at the centro-parietal region, which overlaps with the topographies reported for the cognate effect-related beta desynchronization (**Figure 2b**).

Together, these results confirmed that picture-object processing induces an increase of theta activity across all conditions. Interestingly, and in line with our hypothesis, the results revealed an increase of theta activity for cognate conditions in the picture-naming task *versus* the size-judgment task. The same effect was not observed for non-cognates.

### Theta-beta PAC during visual semantic processing

Once established a significant increase of theta across all conditions, we investigated whether the activation of knowledge (i.e., semantic, lexical and phonological) indexed by beta activity interacted with processes supported by theta activity (i.e., monitoring control) during lexical access. We used cross-frequency coupling analysis to evaluate the functional relationship between theta phase and beta amplitude during picture-object processing in the two tasks. In detail, we determined the participant-specific peaks in theta and beta power from the regions and time-windows of interest identified in the previous analyses, and we used the modulation index (Tort *et al*. 2010; Biau *et al*. 2022) to approximate theta-beta PAC (**Figure 3b and 3c**).

First, we probed whether theta-beta coupling was indeed observed during picture-object processing in all conditions and tasks. To do so, we tested the mean PAC values against zero in the two time-windows of interest, previously determined for the naming-picture and size-judgment tasks, where the cognate effect on beta oscillatory responses was found (**Figure 3b**). Four one-sample *t*-tests revealed a significant theta-beta phase-amplitude coupling during stimulus processing, across all conditions [NamingNC: 0.028 ± 0.004, *t* (17) = 32.23, *p* < .001, Cohen’s *d* = 7.597, BF10 = 3.350e+13; NamingC: 0.029 ± 0.004; *t* (17) = 29.493, *p* < .001, Cohen’s *d* = 6.951, BF10 = 8.285e+12; Size-judgmentNC: 0.032 ± 0.002, *t* (17) = 60.06, *p* < .001, Cohen’s *d* = 14.156, BF10 = 6.354e+17; Size-judgmentC: 0.031 ± 0.003, *t* (17) = 48.259, *p* < .001, Cohen’s *d* = 11.375, BF10 = 1.951e+16; Significant *p*-values were corrected for multiple comparisons at α = .0125]. The ANOVA’s results revealed a significant effect of Task on theta-beta PAC [F (1,17) = 20.345, *p* < .001, η*p2* = .545]. Instead, the main effect of Cognate Status [F (1,17) = 0.009, *p* = .925, η*p2* = .0005] and the interaction between Task and Cognate status [F (1,17) = 2.064, *p = .*169, η*p*2 = .108] were not significant. Finally, as no cognate effect on theta-beta PAC was observed in the picture-naming or the size-judgment task, non-cognate and cognate trials were collapsed together to further explore the difference of theta-beta coupling between the tasks. Results revealed that beta power’s fluctuation concentrated more towards the theta phase during the size-judgment task as compared to the picture-naming task (**Figure 3c**).

To summarise, despite theta-beta interactions were triggered across all conditions and tasks, these interactions were overall stronger during the size-judgment task as compared to the picture-naming task. Interestingly, the cognate status did not modulate the strength of the interaction between theta and beta frequencies.

## 4. Discussion

Recent research work has shown that during object recognition tasks the intention to name an object modulates neural responses reflecting activation of words (Strijkers *et al*. 2012; Branzi *et al*. 2021). However, previous literature has not clarified yet various aspects regarding the nature of this modulation. What are the neural mechanisms underpinning lexicalisation processes? Is lexical access achieved differently depending on the task at hand? The present study answers these questions with some key results, as summarised below.

### Lexical access is enabled by beta desynchronisation independently from the task

First, we established that lexicalisation processes occur even in absence of explicit word-retrieval for oral production. In fact, behavioural and neural cognate effects were observed not only during the picture-naming task, but also during the size-judgment task. Similar results have been observed also by (Strijkers *et al*. 2012), who manipulated word frequency and examined whether ERP responses time-locked to picture-object’s presentation varied depending on whether the task required explicit word-retrieval (picture naming task) or whether it required a semantic judgment but not explicit word-retrieval (semantic categorisation task). Their results revealed a lexical frequency effect irrespective of the intention to name an object. Nevertheless, the interpretation of this effect as being purely lexical was limited by the fact that word frequency tends to correlate with visual and conceptual variables. In other words, it is difficult to establish whether the frequency effect observed by Strijkers *et al*. (2012) was purely lexical, or rather reflected activation of a combination of visual, conceptual, and lexical information. In the present study, we manipulated the cognate status of the stimuli, which is not correlated with any perceptual or conceptual variable (e.g., Costa *et al*. 2000; Costa *et al*. 2005; Christoffels *et al*. 2007; Strijkers *et al*. 2010; Palomar-Garcia *et al*. 2015). Therefore, we can confidently conclude that lexicalisation processes indexed by the cognate effect occur irrespective of the intention to name an object.

Interestingly, the present results contrast with our recent functional magnetic resonance imaging study (Branzi *et al*. 2021). Indeed, we found that explicit word-retrieval in the picture naming task facilitated activation of lexical/phonological representations (measured via cognate effect), by modulating functional connectivity between areas involved in visual object recognition and phonological control. However, this neural cognate effect was not observed in the size-judgment task, which did not require explicit word-retrieval of the object names. The apparent discrepancy between our previous results and the present study might be due to the type of bilinguals tested. In the present study, bilinguals were early and high-proficient in both languages. in our previous study, instead, multilinguals were much less proficient in their non-native third language (i.e., L3) as compared to their L1. Language proficiency might explain these inconsistencies because the cognate effect, which measures the languages’ co-activation, is modulated by the strength of the links between conceptual and lexical/phonological representations (Brenders et al. 2011). Therefore, it is possible that in our previous study (Branzi *et al*. 2021) we did not observe a neural cognate effect in the size-judgment task because the semantic analyses required to perform this task were too superficial to engage the weak links between concepts and L3 lexical/phonological representations. Altogether, the results reviewed above suggest that lexicalisation processes are triggered by visual picture-object processing, independently of the intention to name an object, and the strength of the links between conceptual and lexical/phonological representations might determine the extent to which spreading activation from the semantic system reaches the lexical and phonological representations (especially in the case of multilinguals).

Our findings indicate that the intention to name an object “speeds up” lexical access mechanisms as compared to a non-verbal context. Indeed, picture-object presentation and the cognate effect induced beta desynchronisation at similar centro-parietal scalp regions in both tasks, suggesting that lexical access in the brain is supported by the very same neural mechanism. However, the timing of such cognate-related beta desynchronisation differed depending on the task and took place earlier in the picture-naming task (160 to 260 ms) as compared to the size-judgment task (260 to 380 ms).

Interestingly, Strijkers *et al*. (2012) reported similar time-windows for the neural effect reflecting lexical access in a verbal *versus* non-verbal task comparison. In their study, the ERPs elicited by naming objects with low frequency names started to diverge from those with high frequency names as early as 152 ms after stimulus onset. Instead, during non-verbal categorization, the same frequency effect appeared 200 ms later (∼ 350 ms after stimulus onset). However, regarding the nature of the modulation induced by the intention to speak, the authors reached a conclusion different from ours. Based on the observation that the ERPs measured in these two different time-windows elicited two qualitatively different neural responses in the two tasks (i.e., a P2 and a N400), the authors concluded that lexical access was relying on qualitatively different neural processes. That is, not only the two tasks differed because of the speed with which concepts triggered word activation, but also in the way words were activated.

A plausible alternative to explain the qualitative differences in the ERPs across tasks might reside in differences in the experimental paradigms. In fact, in Strijkers *et al*. (2012) the semantic categorisation task did not differ from the picture-naming task only because it did not require explicit word-retrieval. A key difference was that the semantic categorisation task was a Go/No-go task, and therefore in the semantic categorization task, the analysed trials were no-go trials. Differently from those measured in the picture-naming task (go trials), no-go trials require to withhold a response, a cognitive operation that typically elicits modulation of the N2 and N400, rather than the P2. This interpretation is in line with the finding of a N400 frequency effect for no-go trials (Strijkers, 2012). Consequently, it remains unclear whether the different ERP components observed in the two tasks by Strijkers et al. (2012) reflected different types of lexicalisation processes, differences in the experimental paradigm employed, or a combination of both. In contrast, our study involved two tasks requiring a response for each trial and manipulated a variable that reflects a pure lexical/phonological effect. Our results indicate that lexical/phonological access during object naming and object categorization likely origins from the same spreading activation mechanism, and that intention to speak speeds up lexical access enabled by power decreases in beta oscillations (Amoruso et al. 2021; Geng *et al*. 2022; Piai and Zheng, 2019).

### Monitoring control mechanisms are reflected by theta synchronisation

A second key result refers to the finding that theta (3-7 Hz) increased during picture-object processing, across all conditions and tasks. Further, the topography of such increase peaked at the centro-parietal region. As for the responses observed in the beta band, the fact that theta activity was significantly modulated by picture-object processing in the two tasks, suggests that theta oscillations may support similar cognitive operations in verbal and non-verbal semantic tasks, although with different timings (earlier in the picture-naming as compared to the size-judgment task). This interpretation aligns with previous evidence that established an association between theta rhythms and retrieval of information across different domains, including language and memory (Jensen and Lisman 1998; Bastiaansen et al. 2005; Lisman and Buzsaki 2008). In the language domain, previous studies have reported 4-8 Hz theta power increases and 8-25 Hz alpha– beta power decreases in association to the retrieval of lexical-semantic information from long-term memory (Bastiaansen *et al*. 2005; Piai, Roelofs and Maris 2014; Piai *et al*. 2015; Piai et al. 2016; Piai *et al*. 2017; Geng *et al*. 2022). Furthermore, various electrophysiological studies have employed interference paradigms to investigate the control processes deployed to solve lexical/semantic competition during the picture-name retrieval and showed a link between these processes and theta activity (Piai, Roelofs, Jensen, et al. 2014; Shitova et al. 2017; Krott et al. 2019). The view that theta oscillations synchronise when conflict between representations increases (Cavanagh and Frank 2014; Cohen 2014; Geng *et al*. 2022), accords with our finding of increased theta power during the picture-naming task *versus* size-judgment task for cognates only. In fact, if the intention to name an object induces strong dual-language activation, this should affect especially cognates, because only this category of stimuli receives direct activation from both languages. This dual-language activation should increase conflict between target and non-target cognate words. Since during picture-naming only one of the two languages (the target language) must be selected, as compared to non-cognates, the processing of cognates may require greater monitoring control to avoid erroneous responses (selection of the non-target language). We propose that such increase of monitoring control is reflected by the cognate-related increase of activity in theta oscillatory correlates.

Noteworthily, although dual-language activation occurs also in the size-judgement task, as indicated by the significant cognate effect, such effect seems not only to occur earlier, but also to be stronger in the picture-naming task. Indeed, in the latter, the processing of cognate *versus* non-cognate stimuli resulted in increased accuracy as well as in faster performance. However, in the size-judgment task, the processing of cognate stimuli only increased accuracy as compared to non-cognate stimuli. Therefore, the greater synchronisation of theta oscillations observed for cognates during the picture-naming task *versus* the size-judgment task likely reflects the consequences of stronger dual-language activation induced by the intention to name an object. Despite resulting in facilitatory behavioural performance, dual-language activation might have increased the need of monitoring control processes to ensure that the target language is selected (Li and Gollan 2018).

### The intention to speak modulates theta-beta phase-amplitude coupling during visual object processing

Finally, we explored whether the functional relationship between theta and beta activities depended on the intention to name an object and the cognate status by means of phase-amplitude coupling analysis. Our results revealed first that the coupling at the centro-parietal regions was greater in the size-judgment as compared to the picture-naming task. Second, in both tasks the cognate status did not influence the cooperation between the two oscillatory activities.

The present study may be the first to compare theta-beta phase-coupling directly between a verbal and a non-verbal semantic task using the same stimuli. If the theta-beta coupling reflects a mechanism for the collaborative functioning of lexical activation (beta effects) and response monitoring (theta effects), then although speculative, a plausible interpretation is the increase of theta-beta coupling observed for the size-judgment task might reflect increased monitoring control during the activation of object-related lexico-semantic information (beta activity). Since lexical representations get activated even without the intention to speak, additional control processes might be required to ensure that the correct response (‘yes’ or ‘no’ response in the size-judgment task) is produced as output, rather than the object name itself. Future studies are needed to establish if control demands during language tasks affect neural coupling between theta-beta frequencies.

## Conclusion

The present study provides new evidence for the psycholinguistic models of language production and beyond. The presence of a cognate effect in both our tasks contrasts with concept selection models, which postulate that activation will propagate from concepts to lexical and phonological representations only under the intention to name an object (Levelt, 1989; Bloem and La Heij, 2003). Instead, models incorporating spreading activation from the semantic to the lexical system appear to account better for these data. Accordingly, any activated conceptual representation would lead to activation of lexical information no matter what the goal of the task is (e.g., Caramazza, 1997; Dell 1986; Levelt *et al*. 1999). To account for the different time course of the cognate effect between the two tasks, the spreading of activation principle specified in the models above should consider the importance of contextual aspects and top-down knowledge that proactively impact the interaction between concepts and lexical representations.

Furthermore, our results show that word-retrieval in verbal and non-verbal semantic tasks rely on similar oscillatory dynamics for response selection and activation of words, a finding which aids the current debate on the domain-specificity of neural processes supporting language core-operations (Fedorenko and Shain 2021). In doing so, here we demonstrate that a core language operation, such as lexical access, may occur in absence of oral production. This finding has important implications for understanding the performance of both healthy individuals and neurological patients in verbal and non-verbal semantic tasks.

## Supplementary Information

This Supplementary file includes:

Cognate effect (non-cognates *versus* cognates) modulation on alpha (8-12 Hz) oscillations during the picture-naming and size-judgment tasks.

We addressed whether mean alpha (right) power increased during the picture-naming and size-judgment tasks, as compared to the pre-stimulus baseline (**Figure S1**). Four independent one-sampled *t*-tests (two-tailed) revealed no significant increase (or decrease) of alpha activity during stimulus processing in all the conditions (NamingNC: 0.06 ± 0.28, *t* (17) = 0.94; *p* > .5, Cohen’s *d* = 0.22; NamingC: 0.03 ± 0.27, *t* (17) = 0.47, *p* > .5, Cohen’s *d* = 0.11; SemanticNC: -0.14 ± 0.34, *t* (17) = -1.78, *p* = 0.37, Cohen’s *d* = -0.42; SemanticC: -0.19 ± 0.33, *t* (17) = -2.47, *p* = .1, Cohen’s *d* = -0.58; Significant *p*-values were corrected for multiple comparisons at α = .0125). A two-way repeated-measure ANOVA with the factors Task (picture-naming vs. size-judgment) and Cognate Status (non-cognate vs. cognate) revealed a significant effect of Task on mean alpha power [F (1,17) = 11.11, *p* < .001, η*p*2 = .40]. No effect of Cognate status [F (1,17) = 1.80, *p* = .20, η*p*2 = 0.1] or interaction between Task and Cognate status [F (1,17) = 0.05, *p* = .83, η*p*2 < .001] was found.

**Figure S1.**
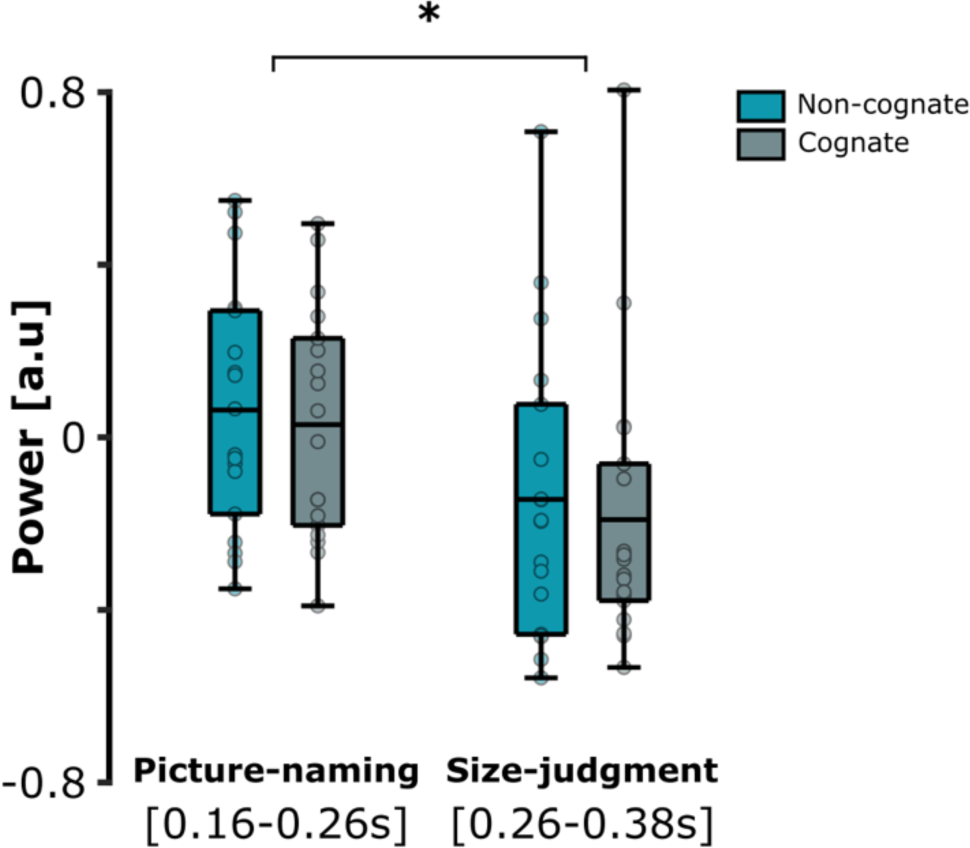
Boxplots of the mean alpha power (8-12 Hz) across the cognate and non-cognate conditions and tasks. The time-window of interest for the naming and size-judgment task were respectively 0.16 to 0.26 s and 0.26 to 0.38 s after the onset of stimulus presentation (significant differences evidenced with black stars). The bottom part of the boxplots indicates the first quarter of the score distributions, the upper part of the boxplots indicates the third quarter of the score distributions. The error bars indicate the minimum and maximum scores from the distributions.

### Cognate effect in the picture-naming and size-judgment tasks with the same pool of electrodes

We tested whether the main result reported in the manuscript (i.e., cognate effect reflected by beta desynchronization, occurring in distinct time-windows for the picture-naming and the size-judgment tasks), was not driven by the difference of electrode pools included in the analysis. To this end, we re-run the exact same analysis (non-cognate *versus* cognate) on the time-frequency decomposition of the spectral power difference in the two tasks, but with a common pool of overlapping electrodes. In this control analysis, for both tasks the cluster of interest contained the following electrodes: C3, CP3, P3, C4, CP4, P4, Cz, CPz, and Pz (**Figure S2**). The normalised mean beta power was computed for all conditions across the common cluster in the two time-windows of interest (picture-naming: 160-260 ms; size-judgment: 260-380 ms with respect to the stimulus onset). The four independent one-sample *t*-tests (two-tailed) confirmed that the mean beta power was significantly below zero as compared to the baseline preceding the stimulus onset, across all conditions: NamingNC: -0.14 ± 0.15, *t* (17) = -3.79, *p* = .001, Cohen’s *d* = -0.89; NamingC: -0.23 ± 0.1; *t* (17) = -10, *p* < .001, Cohen’s *d* = -2.4; Size-judgmentNC: -0.14 ± 0.15, *t* (17) = -3.87, *p* = .001, Cohen’s *d* = -0.91; Size-judgmentC: -0.21 ± 0.15, *t* (17) = -6.02, *p* < .001, Cohen’s *d* = -1.42; Significant *p*-values were corrected for multiple comparisons at α = .0125). The ANOVA’s result confirmed a significant effect of Cognate status [F (1,17) = 13.67; *p* = .002, η*p2* = .45], establishing a greater beta desynchronisation for cognates as compared to non-cognates. No significant main effect of Task [F (1,17) = 0.01, *p* = .91, η*p*2 < .001] or interaction between Task and Cognate status [F (1,17) = 0.24, *p* = .63, η*p*2 = .01] was found. Therefore, picture-object processing induced a significant beta desynchronisation independently from the task and condition, in line with the results reported in the main text (Figure 2). Further, the cognate stimuli induced a greater beta desynchronisation of the beta oscillations as compared to non-cognate stimuli in both tasks. Altogether, this analysis validates the results reported in the main manuscript, even when performed on a common cluster overlapping with the two separate regions of interests between the picture-naming and size-judgment tasks.

**Figure S2.**
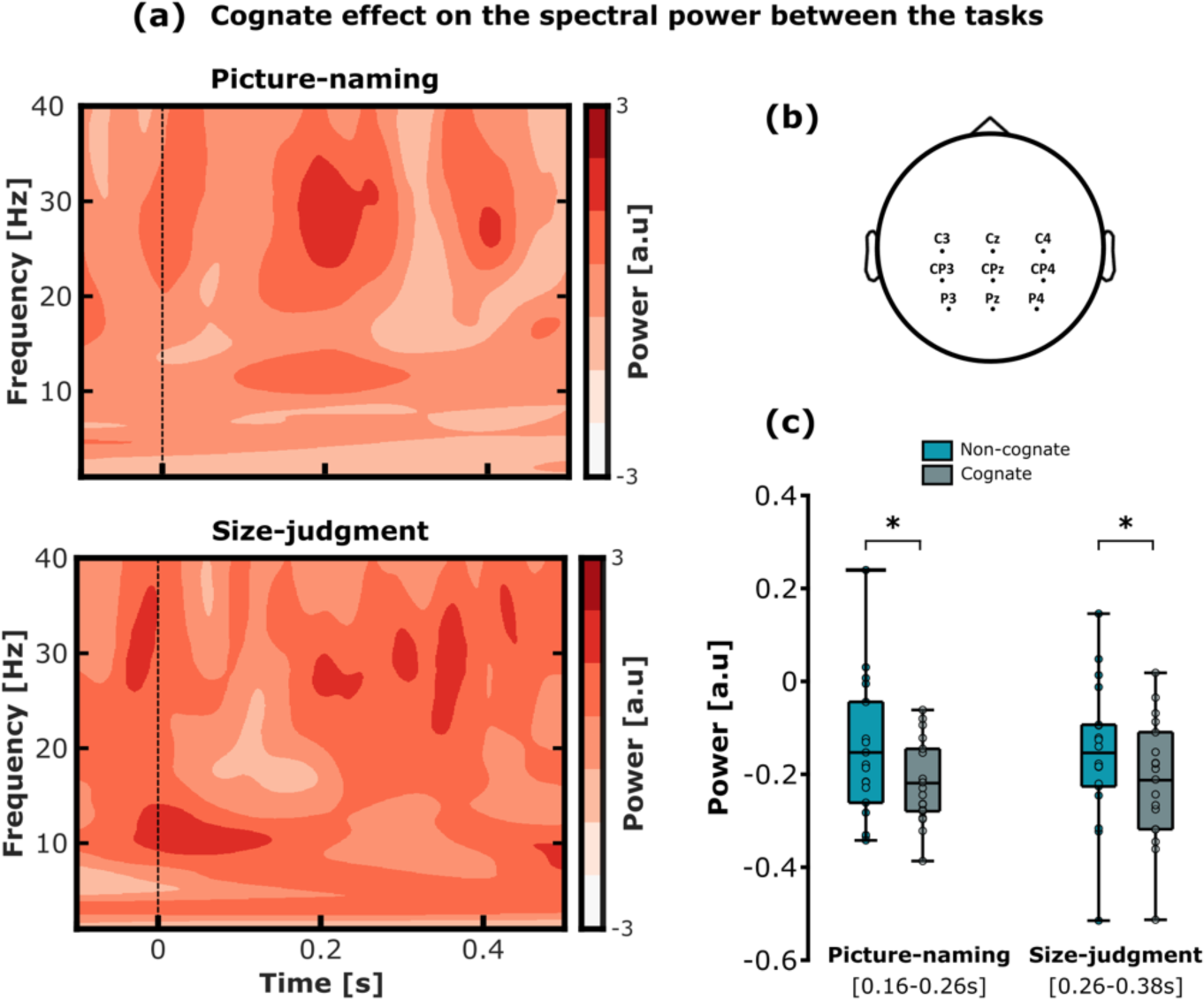
Cognate effect (non-cognates *versus* cognates) modulation on beta oscillations during the picture-naming and size-judgment tasks, using the same pool of electrodes. (a) Time-frequency representations (TFRs) of the spectral power difference, i.e., non-cognate *versus* cognate, in the picture-naming (upper part) and in the size-judgment task (bottom part). The TFRs represents the average of all electrodes included in the common cluster depicted in the topoplot of panel b. (b) Pool of electrodes constituting the common cluster used in the control analysis: C3, CP3, P3, C4, CP4, P4, Cz, CPz, and Pz. (c) Boxplots of the mean beta power (25-35 Hz) across the cognate and non-cognate conditions and tasks (significant differences evidenced with black stars). The time-window of interest for the naming and size-judgment task were identical to the analysis reported in the main manuscript, i.e., 160 to 260 ms and 260 to 380 ms after the stimulus presentation onset, respectively. The bottom part of the boxplots indicates the first quarter of the score distributions, the upper part of the boxplots indicates the third quarter of the score distributions. The error bars indicate the minimum and maximum scores from the distributions.

### Searchlight-based analysis assessing the magnitude of the cognate effect on beta power over the scalp

We performed a searchlight-based analysis to contrast the amplitude of the cognate effect on beta power (i.e., non-cognate-cognate) in the significant clusters of interest against all the remaining electrodes of the scalp (**Figure S3**). This analysis revealed that the cognate effect modulation on beta power in the regions of interest was significantly greater than any other size-matching searchlight-based regions over the scalp (*p* < .001 in both the picture-naming and size-judgment tasks).

**Figure S3.**
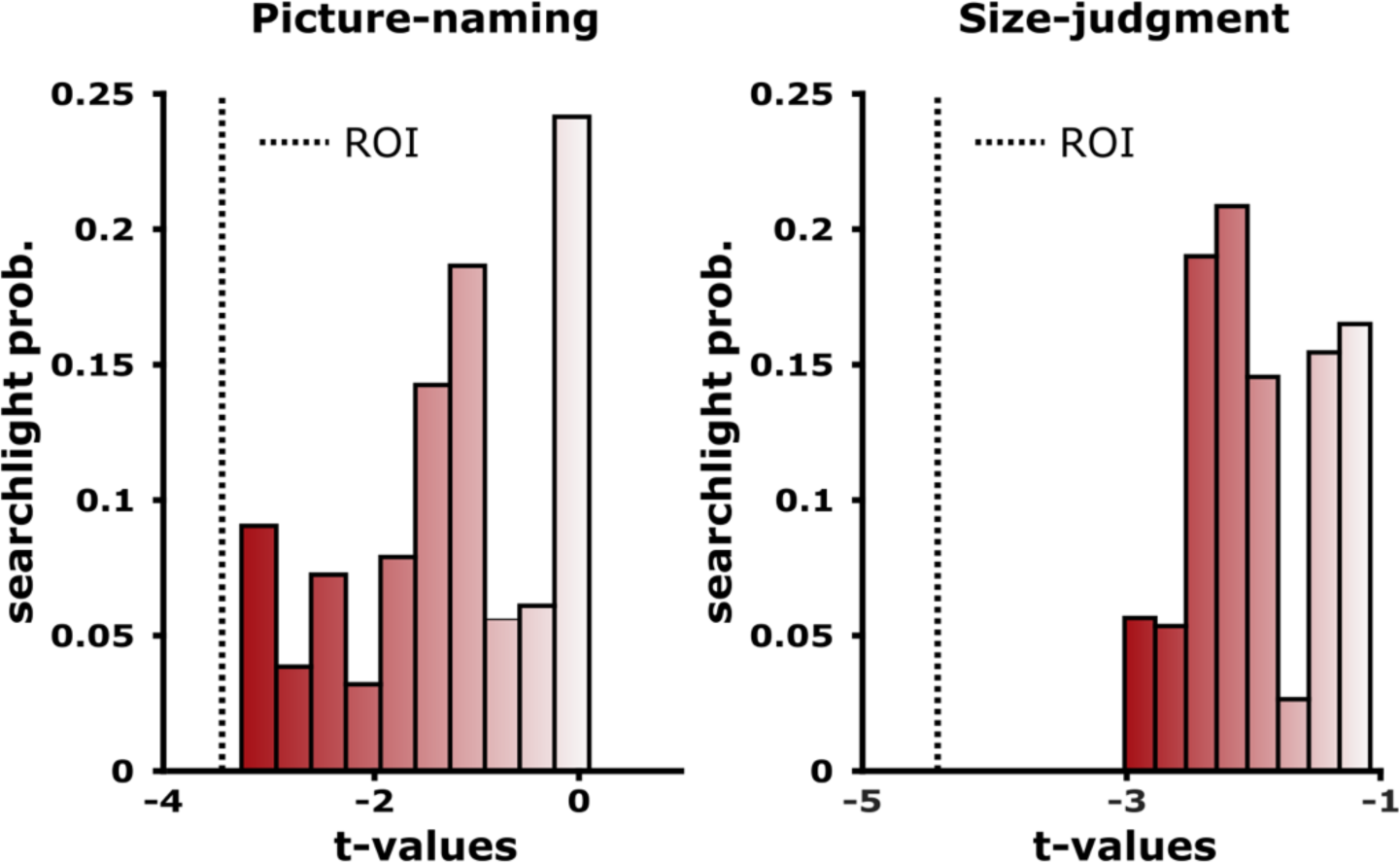
Distribution of *t*-statistics for the cognate effect on beta power in the regions of interest for every searchlight over the scalp, in the two tasks. The absolute *t*-value for the regions of interest (non-cognate *versus* cognate; *t*-values of the ROIs evidenced by the dashed line) was greater than the random searchlight-based mini-clusters matching the size of the regions of interest for both the picture-naming and size-judgment tasks (total number of permutations = 2000; *p*-values < .001).

### Correlation between beta and theta power and behavioural accuracy in the picture-naming and size-judgment tasks

In a set of exploratory analyses, we examined whether the decrease in beta band observed for the cognate effect (non-cognates *versus* non-cognates) predicted differences in accuracy between non-cognate and cognate conditions. We correlated (Pearson’s correlation) the non-cognate *versus* cognate difference of normalized mean beta power (25-35 Hz) from the regions of interest [Δpower = beta power non-cognate - beta power cognate] with the non-cognate *versus* cognate difference of correct responses [Δcr = CRnon-cognate - CRcognate] in the picture-naming and size-judgment tasks separately (**Figure S4**). For each participant, the mean beta power (25-35 Hz) was computed in the four conditions with the same procedure reported in the main manuscript. The results revealed no significant correlation between the correct response rate differences (Δcr) and the amplitude of beta power differences (Δpower) in the picture-naming (*r* = 0.07, *p* = .78; two-tailed) or the size-judgment task (*r* = 0.19, *p* = .46; two-tailed). Noteworthily, the behavioural performance (accuracy and response times-RTs) were not significantly correlated between the picture-naming and size-judgment task, suggesting that a similar cognate effect on performances was not driven by individual patterns (see **Figure S5b**).

**Figure S4.**
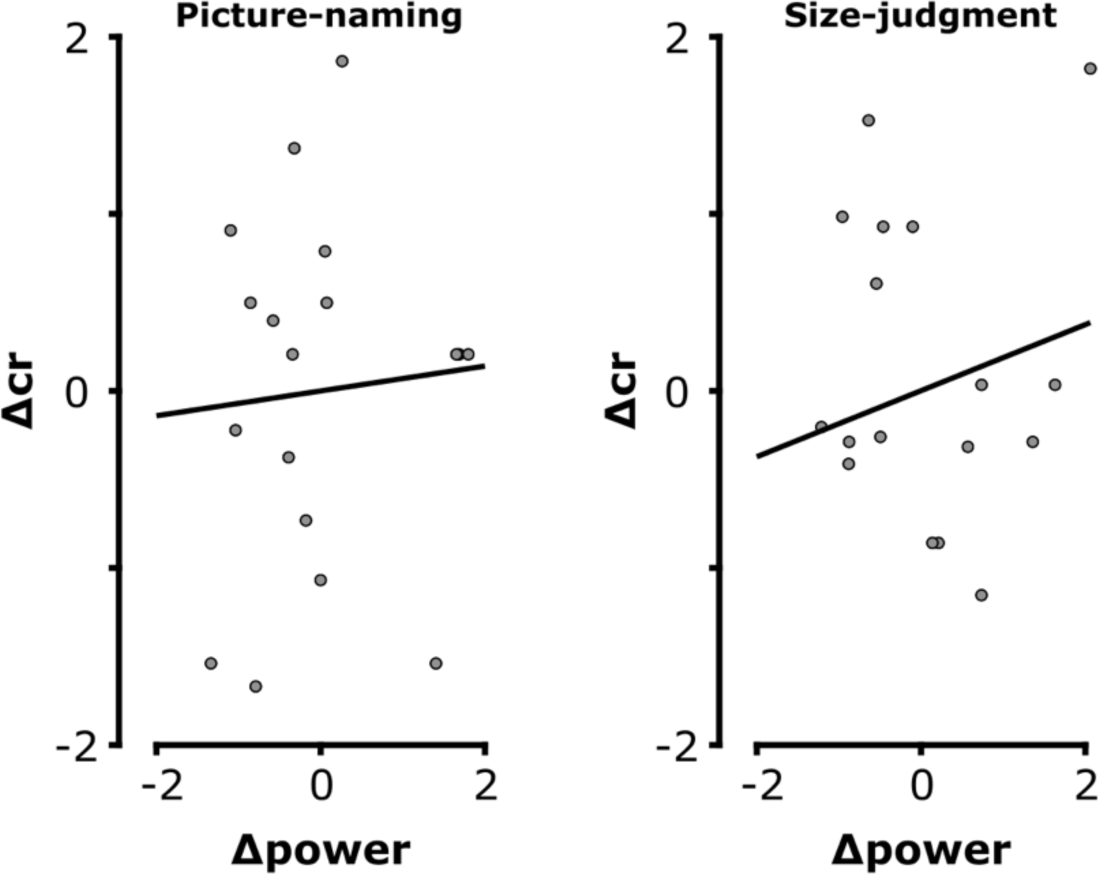
Correlations between behavioural (accuracy) and neural (beta power) cognate effect measured in the picture-naming and the size-judgment tasks separately. The difference of beta power between the non-cognate *versus* cognate condition (Δpower; *x*-axis) did not significantly correlate with the difference of accuracy between the non-cognate *versus* cognate condition (Δcr; *y*-axis) in the picture-naming task (left scatterplot) or in the size-judgment task (right scatterplot).

We also examined whether the increase in theta band found for cognates in picture naming *versus* size judgment task predicted differences in accuracy for the same contrast. Therefore, for cognates only we correlated (Pearson’s correlation) the between-tasks difference (picture naming *versus* size-judgment task) of normalized mean theta power from the regions of interest [Δpower = theta power cognate picture naming - theta power cognate size-judgment] with the between-tasks difference of correct responses [Δcr = CR cognate picture naming – CR cognate size-judgment]. For each participant, the mean theta power (3-7 Hz) was computed in the four conditions with the same procedure reported in the main manuscript. The results revealed no significant correlation between the correct response rate differences (Δcr) and the amplitude of theta power differences (Δpower) for cognate stimuli between tasks (*r* = 0.068, *p* = .7862; two-tailed).

### Picture-naming and size-judgment tasks: between-tasks spectral power differences and between -tasks correlations of behavioural performance

We examined whether the timing difference relative to the cognate effect (beta desynchronization) observed between the two tasks was driven by differences in the type of (verbal *versus* manual) response preparation processes. To do so, we investigated the difference of power spectrum between the picture-naming and size-judgment tasks, by collapsing the cognate and non-cognate trials together. The resulting TFR shows that there was no difference of power in the low frequencies of interest (< 35 Hz) within the time-window where the cognate effect was found (i.e., approximatively from 100 to 400 ms with respect to the stimulus onset) (**Figure S5a**). The cluster-based permutation tests revealed a significant positive cluster (*p* = .05, cluster size = 1.08e4) and a significant negative cluster (*p* < .001, cluster size = -4.36e4). In detail, TFRs revealed a task (picture naming *versus* size-judgment) difference in the alpha-beta band (∼ from 10 to 25 Hz) around 400 ms after the picture onset. Since in the picture naming task the cognate effect was observed between 160-260 ms after stimulus onset, it is unlikely that task-differences in the type of response (observed around 400 ms after stimulus onset) account for the results reported in our study.

**Figure S5.**
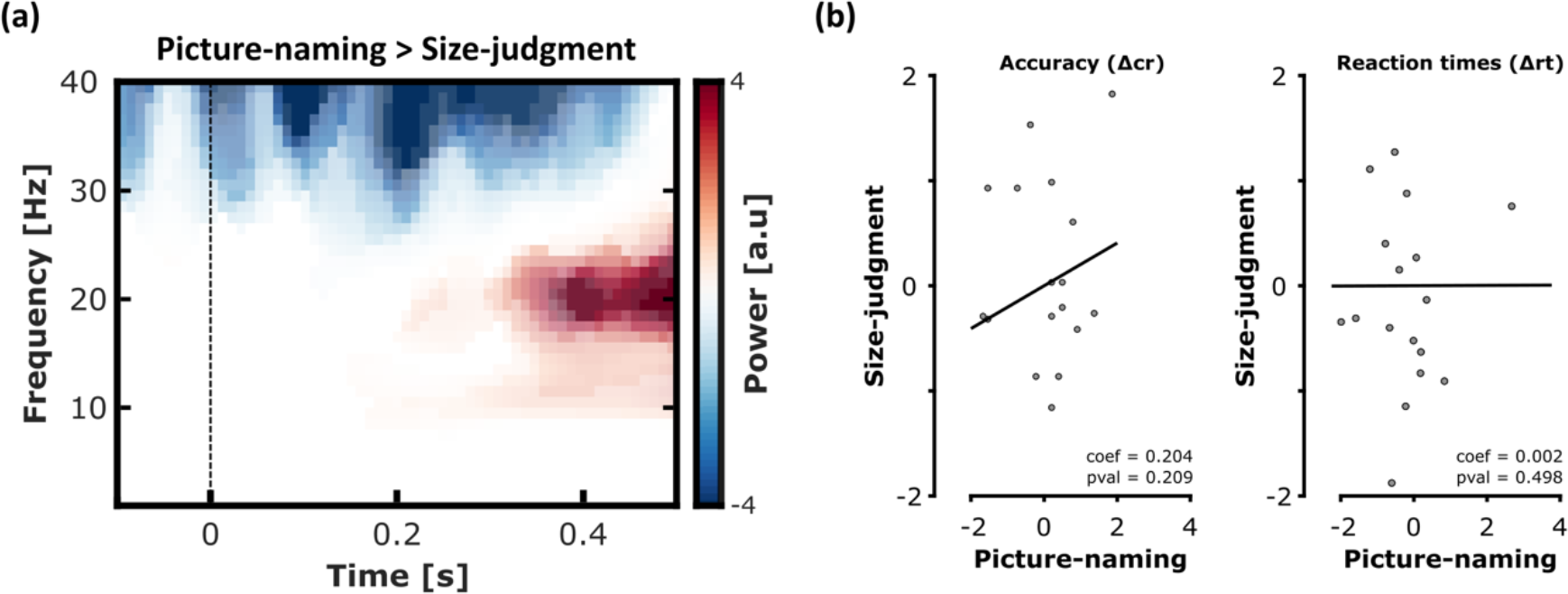
Spectral difference of power between the picture-naming *versus* size-judgment task (and viceversa), and correlation between behavioural cognate effect in the two tasks. (a) The TFR represents the average of electrodes included in the common cluster used in the control analysis (C3, CP3, P3, C4, CP4, P4, Cz, CPz, and Pz). (b) Left scatterplot: the difference of accuracy between the non-cognate *versus* cognate condition (Δcr) in the picture-naming task (*x*- axis) did not significantly correlate with the difference of accuracy between the non-cognate *versus* cognate condition in the size-judgment task (*y*-axis). (b) Right scatterplot: the difference of RTs between the non-cognate *versus* cognate condition (Δrt) in the picture-naming task (*x*-axis) did not significantly correlate with the difference of RTs between the non-cognate *versus* cognate condition in the size-judgment task (*y*-axis).

Finally, we examined the relationship between the behavioural cognate effect measured in the picture-naming and the size-judgment tasks. We computed the cognate effect for correct responses [Δcr = CRnon-cognate - CRcognate] as well as for RTs [Δrt = RTnon-cognate - RTcognate] in both tasks separately (**Figure S5b**). Then we correlated the behavioural cognate effects (both accuracy and RTs) between tasks, using two separate Pearson’s correlation analyses. Results revealed no significant correlation between the cognate effects in the two tasks [accuracy (ΔCR): *r* = 0.20; *p* = .21; two-tailed; RT (ΔRT): (*r* = 0.002, *p* = 0.498; two-tailed).

## Acknowledgments

This work was supported by a Sir Henry Wellcome Postdoctoral Fellowship awarded to E.B. (Grant reference number: 210924/Z/18/Z). C.D.M received funding from the European Research Council (ERC) under the European Union’s Horizon 2020 research and innovation programme (Grant Agreement No: 819093) and the Spanish Ministry of Economy and Competitiveness (PID2020-113926GB-I00). The authors acknowledge the Basque Government through the BERC 2022-2025 program and by the Spanish State Research Agency through BCBL Severo Ochoa excellence accreditation CEX2020-001010-S.

